# Marriage in the Melting Pot: An evolutionary approach to European ancestry, homogamy and fertility in the United States

**DOI:** 10.1101/2020.05.25.114108

**Authors:** Alexander Schahbasi, Susanne Huber, Martin Fieder

## Abstract

**Objective:** To understand marriage patterns, homogamy and fertility of women of European ancestry in the United States from an evolutionary perspective we aim to investigate if a prevalence for ancestral homogamy exists, the factors influencing a female preference for an ancestral homogamous vs. an heterogamous marriage, if an ancestral homogamous vs. heterogamous marriages influences fertility and if there is an inherted component of the tendency to marry homogamously vs. heterogamously. Furthermore we aim to determine the heritability of homogamous vs. heterogamous marriage behaviour.

**Methods:** We used the census data of 369,121 US women married only once and aged between 46 and 60 years, provided by IPUMS USA (https://usa.ipums.org/usa/). We used linear mixed models to determine associations of the probability of a homogamous vs. heterogamous marriage and the individual fertility of a women. We aimed to estimate the heritability (in our case genetic & parental environment) of marriage behaviour using a linear mixed model.

**Results:** We found, that ancestral heterogamous marriages are more frequent (56.5%), compared to homogamous marriages (43.5%). Most of the variance in inter- ancestry marriage and fertility is explained by ancestry per se, followed by the ratio of individuals of a certain ancestral background in a county: the more individuals of a certain ancestry live in a county the lower is the tendency to marry someone of a different ancestral background. Furthermore we found that about 11.8% of the marriage behaviour is heritable. Being in a homogamous marriage as well as the income of the spouse are both significantly positively associated with the number of children a women has and the probability that a women has at least one child.

**Discussion:** The most important explaining factor (in terms of variance explained) for being in an ancestral homogamous vs. heterogamous marriage, for number of children, as well as childlessness is the ancestry of the women. Albeit we are not able to distinguish the genetic and social heritability on basis of our data, with a total value of 11.8% variance explained, only a small heritability for in-group vs, out-group marriage behaviour is indicated.

## Introduction

Investigating marriage patterns, homogamy and fertility involves various perspectives that have to be taken into consideration, particularly with regards to the long-term implications of migration. Since the findings of David Reich (Reich 2018) the enormous genetic impact of migration and admixture in shaping the human genome has become evident. These findings on the global dispersal of Homo Sapiens allow for the classification of migration processes and subsequent admixture as an inherent trait of the human species. However, these migration flows induced processes of human interactions taking place on a continuous spectrum ranging from intermarriage/admixture to hostile/violent encounters at its extremes. Understanding these extremes helps to comprehend long-term social developments in the context of migration and social cohesion.

Beginning on one side of the spectrum, looking at violent encounters it has to be noted that violence declined throughout the course of human history. Steven Pinker (Pinker 2011) has documented this process and attributed this development to the Leviathan (the state), trade, feminization, globalization and reason. In addition, Richard Wrangham (Wrangham 2019) has argued that Homo sapiens has undergone a process of self-domestication (particularly with regards to reactive violence). We have previously argued that within societies that are exposed to migration there are always individuals who are more open to migration (and exchange) while other remain wary or hostile (Schahbasi et al. 2020 in press). It can be assumed that these response patterns have been shaped in our evolutionary history and that these mechanisms are still in place today. Both attitudes made evolutionary sense for communities to thrive: On the on hand there was a necessity to avoid inbreeding (Sikora et al. 2017) and allow for cultural exchange, but on the other hand there was a necessity to defend against potential lethal interactions.

When considering peaceful social exchanges, it has to be distinguished between interactions with genetically related indivuals and non-geneticall related individuals. Based on the theory of kin-selection (Hamilton 1964) the propensity to cooperate is related to the degree of genetic relatedness. The level of cooperation is thus highest between close relatives (e.g. parents and their children or siblings) and declines with decreasing genetic relatedness. People in our evolutionary past mainly lived in small groups of not more than 150 persons (Dunbar 1993), most of whom were related to some degree. With the onset of the agricultural revolution 10,000 years ago and the settling of people into larger agglomerations (Harari 2015), different modes of cooperation evolved. As cooperation in larger settlements occurs mainly between genetically not related individuals, these are based on the principle of reciprocal altruism (Trivers 1971). These exchanges are based on the principal of reciprocal exchange: to give something expecting a return of some kind. There is of course an overlap between kin selection (inclusive fitness) and recicprocal altruism, but it has been demonstrated that for the latter there is a necessity for punishment (“altruistic punishment”) in case of rule violation (Fehr & Gächter 2002).

In the context of migration and cohabitation of culturally and religiously diverse communities in contemporary urban centers it could be argued that to ensure long-term social cohesion through the highest possible of level of cooperation between groups, there need to be genetic bonds between groups. At the same time, there is a prevailance of homogamy that has been demonstrated for several factors: e.g., body height (Stulp et al 2017, Stulp et al. 2013), religion (reviewed in Fieder & Huber 2016), political attitudes and ethnicity (Blackwell & Lichter 2004, Fu, X., & Heaton 2008). While individuals on the political left and right prefer someone with a similar political attitude, both sides of the political spectrum show a preference for a partner with the same ethnic background - more so on the political right than the political left (Anderson et al 2014). Politically assorted mating is so far among the strongest of all investigated traits (personality, social and biometric) (Alford et al. 2011). Furthermore, educational attainment is an increasingly important factor in the context of mating, particualry for the lower and higher educated strata of a society (Blackwell & Lichter 2004, Fieder & Huber 2011).

From an evolutuionary point of view the questions is, whether assortative mating along certain traits leads to any selective advantages, i. e. whether assortaive mating along a certain trait leads to an increase in the number of children, hence to a fitness gain. Assortative mating and the prevalence of homogamy has been often documented but a potential correlation between homogamy and reproduction has been only sparsely investigated so far. Studies find that educational homogamy is particulary associated with a lower prevalence of childlessness (Huber & Fieder 2011, Bavel 2012, Huber & Fieder 2016) and religious homogamy is positively associated with both fertility and having at least one child (Fieder & Huber 2016). Moreover, religious homogamy may compensate for ethnic heterogamy in terms of reproduction and vice versa (Huber & Fieder 2018). Additionaly, increasing height difference between spouses may increase the risk of a cesarion section (Stulp et a. 2011) and thus may also represent a selective force in the past.

It has been theorized that humans may detect genetic similarities on basis of phenotypic traits and mate assortatively on these traits and that such a behavior may enhance fitness (Rushton et al. 1985, Salter 2009). Marrying somone who is genetically closer may bring reproductive benefits as has been demonstrated on the basis of data from Iceland: the average number of offspring decreases with genetic relatedness from second-order cousins on (Helgason et al. 2008). Thus, during our evolutionary past moderate inbreeding may have led to reproductive benefits (Fox 2015) while leading to a higher homozygosity and an association with health, intellectual and other related problems (Clark et al. 2019).

Based on these deliberations we aim to investigate on basis of US census data i) if a prevalence for ancestral homogamy exists; ii) which factors influence a female preference for an ancestral homogamous vs. an heterogamous marriage; iii) if ancestral homogamous vs. heterogamous marriages influence fertility (measured as number of children and childlessness); and iv) the inherted component of the tendency to marry homogamously vs. heterogamously.

## Methods

To investigate the prevalence of homogamy, the number of children and childlessness according to the ancestry of US women and their spouses we used the following data sets i) ***data set 1***: the census records of 369,121 women married only once aged between 46 and 60 years (almost completed or completed reproduction) and their spouses and ii) ***data set 2***: the census records of 2,721 women aged between 16 and 35 years, married only once, their spouses as well as their parents, all living in the same household. Both data sets have been extracted from the US census of 1980, provided by IPUMS USA (https://usa.ipums.org/usa/) (Ruggles et al. 2020). We only included individuals of European ancestry (Table S2) in our analysis. We further used data set 2 to investigate the proportion of the additive genetic heritability and the proportion of the paternal social heritability of marrying homogamous vs. heterogamous according to ancestry, using a linear mixed model. We included the following variables in the analyses: woman’s age, her age at first marriage, woman’s own education (encoded in 21 steps, further used as continuous variable - supplement Table 1), total yearly income of the woman in a year as well as of her spouse, the ratio of persons of a certain ancestry in a US county (“ratio ancestry county”), number of children born to a woman and her childlessness (encoded: 0 = childless, 1 = one or more children), if a women is in an ancestral homogamous or heterogamous marriage (0 = heterogamous and 1 = homogamous, HHM), and ancestry of the women (encoded as described in supplement Table 2) as well as for data set 2 only, if the parents of a women is ancestral homogamously vs. heterogamously married (0 = heterogamous and 1 = homogamous, PHHM).

### HHM, number of children and childlessness

On basis of data set 1, we calculated the following three separate general linear mixed models: i) age, age at first marriage, education, total income, total income of spouse, and ratio ancestry county, regressing on HHM on basis of a binomial error structure; ii) HHM, age, age at first marriage, education, total income, total income spouse, and ratio ancestry county regressing on number of children on basis of a Poisson error structure, and iii) HHM, age, age at first marriage, education, total income, total income spouse, ratio ancestry county regressing on childlessness on basis of a binomial error structure. In all three models, ancestry of a woman was included as random factor. Mixed models were calculated in R, library MASS, function glmmPQL.

### Genetic and social heritability

On basis of data set 2 we calculated the following linear mixed model (https://cran.r-project.org/web/packages/QGglmm/vignettes/QGglmmHowTo.pdf; de Villemereuil et al. 2016): HHM regressing on the random factor PHHM (redundantly used in the model for both mother and father as 1 if PHHM is homogamous and 0 if PHHM is heterogamous), controlling for age and education as fixed factors, on basis of a binomial error structure (i. e. *HHM = age + education + random (PHHM father) + random (PHHM mother*). We calculated the general linear mixed model using R, library lme4, function glmer. “Heritability” was calculated by the function QGparams from the R library QGglmm.

## Results

### HHM

We found that ancestral heterogamous marriages are more frequent (56.5%) compared to homogamous marriages (43.5%) (Table 1).

**Table 1.** Frequency of HHM.

| **HHM** |  |  |  |
| --- | --- | --- | --- |
|  |  | **Frequency** | **Percent** |
| Valid | heterogamous | 208735 | 56.5 |
|  | homogamous | 160386 | 43.5 |
|  | Total | 369121 | 100.0 |

We further found that age and the ratio of individuals of a certain ancestry in a county are significantly positively associated with being in a homogamous marriage, whereas age at first marriage, spouse’s income and education are significantly negatively associated with being in a homogamous marriage. Most of the variance (31%) in HHM is explained by the ancestry of women (random factor), hence the ancestral background largely determines HHM. From the fixed factors, most of the variance is explained by the ratio of individuals of a certain ancestral background in a county (6%): the more individuals of a certain ancestry live in a county, the lower is the tendency to marry someone of a different ancestral background. Second highest proportion (1.3%) of the fixed factors in the variance of HHM is explained by highest education (Table 2).

**Table 2.** General linear mixed model of age, age at first marriage, education, income, income of the spouse, the ratio of the own ancestry group in a county, regressing on HHM (endcoded as 0 = heterogamous and 1 = homogamous) on basis of a binomial error structure, with ancestry as random factor.

|  | **Beta**<br>**Values** | **Std.Error** | **t-value** | **p-value** | **%**<br>**Variance** |
| --- | --- | --- | --- | --- | --- |
| (Intercept) | 0.000 | 0.162 | -4.814 | P < 0.0001 |  |
| age | 0.020 | 0.001 | 2.806 | 0.005 | 0.021 |
| age marriage | -0.029 | 0.001 | -3.879 | 0.000 | 0.070 |
| education | -0.434 | 0.002 | -53.114 | P < 0.0001 | 1.255 |
| total income | 0.006 | 0.000 | 0.777 | 0.437 | 0.056 |
| total income spouse | -0.090 | 0.000 | -11.660 | P < 0.0001 | 0.334 |
| ratio ancestry<br>county | 1.033 | 0.073 | 63.923 | P < 0.0001 | 5.968 |
| DF | 366452.000 |  |  |  |  |
| (Intercept) | Residual |  |  |  |  |
| StdDev: 1.111785 | 1.000 |  |  |  |  |
| Variance expl.<br>random | 31.4 |  |  |  |  |

### Number of children

We found that age, age at first marriage, women’s own income as well as the ratio of the own ancestry group in a county are significantly negatively associated with a woman’s number of children. Being in a homogamous marriage as well as the income of the spouse are significantly positively associated with her number of children. Education has no significant association with the number of children. Again, the highest proportion of variance is explained by the random factor “ancestry” (12%). Out of the the fixed factors, age of first marriage explains most of the variance (8.6%) followed by a women’s own income (1%). HHM and the ratio of the own ancestry group in a county explain less than 0.1% of the variance.

**Table 3.**
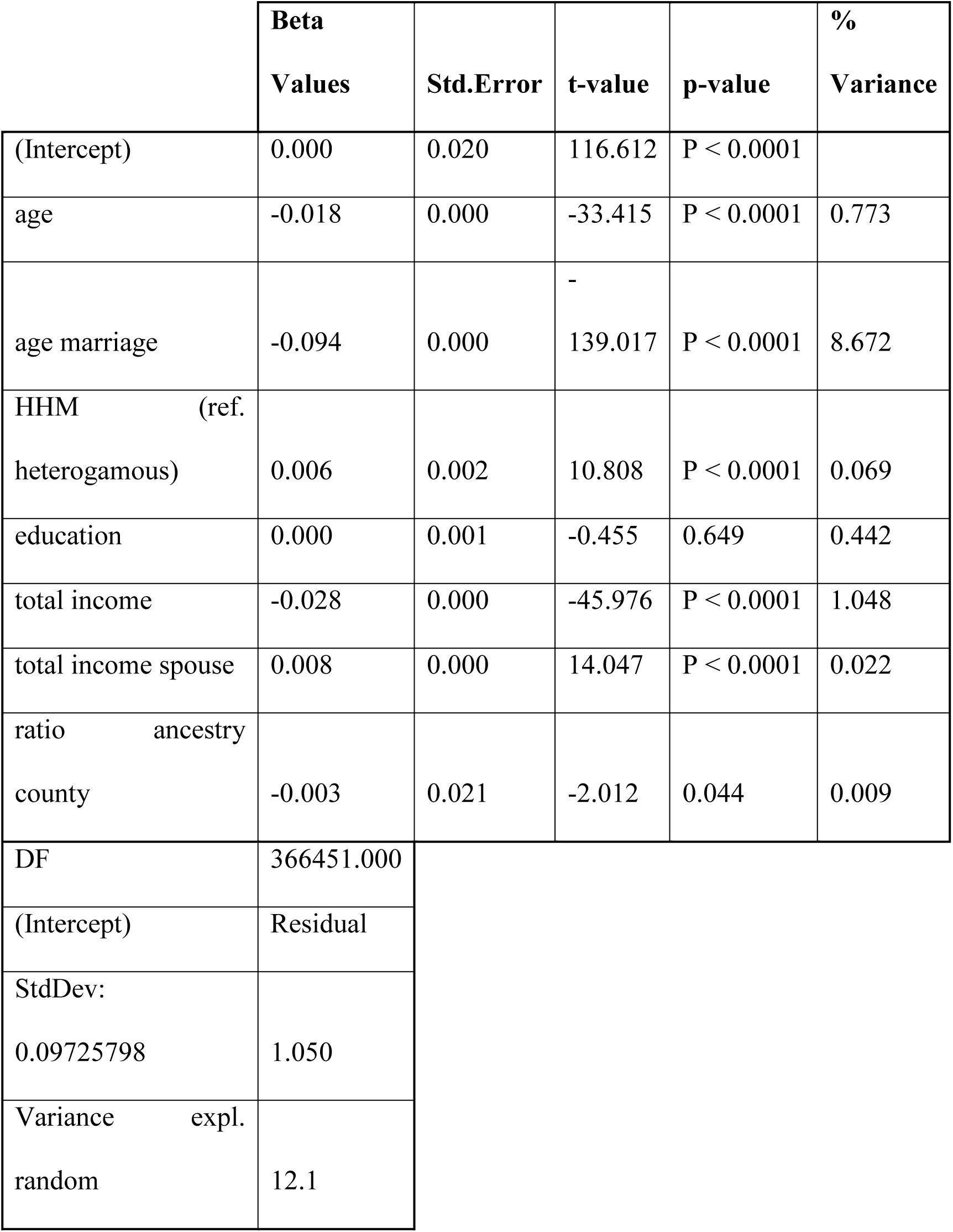
General linear mixed model of age, age at first marriage, education, income, income of the spouse, the ratio of the own ancestry group in a county and HHM, regressing on number of children on the basis of a Poisson error structure, with ancestry as random factor.

### Childlessness

Age, age at first marriage, and women’s own income are significantly negatively associated, whereas being in a homogamous marriage, education as well as the income of the spouse are significantly positively associated with having at least one child. The ratio of the own ancestry group in a county has no significant effect. As is the case in the other models, most of the variance is explained by the random factor “ancestry” (14%). Out of the fixed factors, age of first marriage explains most of the variance (12.4%) followed by a women’s own (~1%) as well as husbands income (~0.9%). Again, HHM and the ratio of the own ancestry group in a county explain less than 0.1% of the variance.

**Table 4.** General linear mixed model of HHM, age, age at first marriage, education, income, income of the spouse, and the ratio of the own ancestry group in a county regressing on childlessness (encoded as 0 = childless, 1 = one or more children) on basis of a binomial error structure, with ancestry as random factor.

|  | **Standardized**<br>**Values** | **Std.Error** | **t-value** | **p-value** | **%**<br>**Variance** |
| --- | --- | --- | --- | --- | --- |
| (Intercept) | 0.000 | 0.091 | 69.316 | P < 0.0001 |  |
| age | -0.236 | 0.002 | -9.828 | P < 0.0001 | 0.873 |
| age marriage | -2.485 | 0.001 | -138.869 | P < 0.0001 | 12.422 |
| ancestry same (ref.<br>no) | 0.177 | 0.014 | 7.081 | P < 0.0001 | 0.090 |
| education | 0.157 | 0.003 | 6.169 | P < 0.0001 | 0.128 |
| total income | -0.593 | 0.000 | -28.695 | P < 0.0001 | 0.971 |
| total income spouse | 0.600 | 0.000 | 21.995 | P < 0.0001 | 0.868 |
| ratio ancestry<br>county | 0.057 | 0.122 | 1.148 | 0.251 | 0.066 |
| DF | 366451.000 |  |  |  |  |
| (Intercept) | Residual |  |  |  |  |
| StdDev: 0.1101647 | 0.998 |  |  |  |  |
| Variance expl.<br>random | 14.1 |  |  |  |  |

### Genetic and Social Heritability of HHM

Regressing HHM on PHHM by using a general linear mixed model with PHHM as random factor and age and education as fixed factors, we found that genetic and social inheritance account for 11.18% of the variance in HHM. Our model thus suggests that 11.18% of the marital behaviour in terms of marrying someone of the same or different ancestry is explained by genetic and social inheritance from the parents.

**Table 5.**
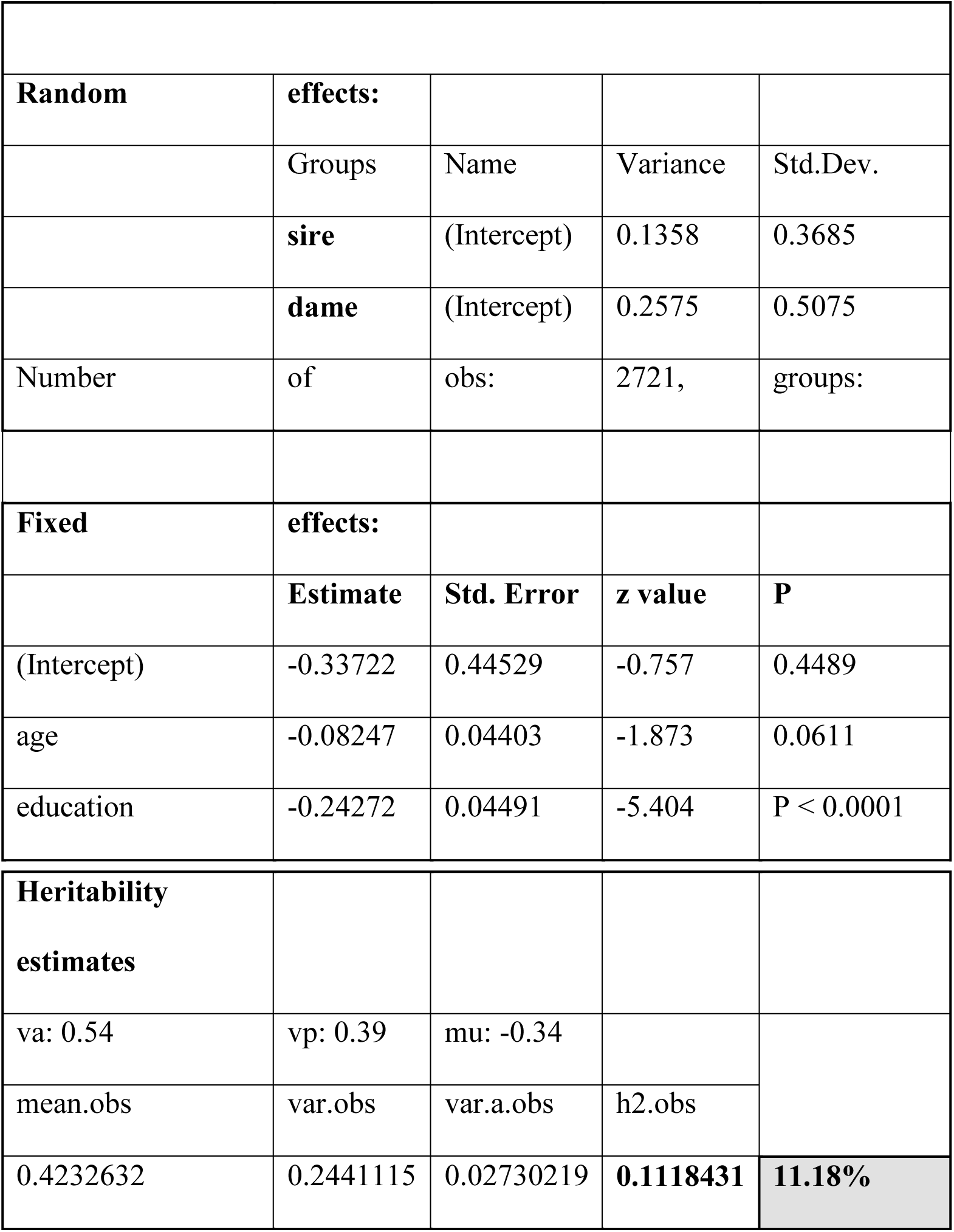
Estimates of social and genetic heritability by a general linear mixed model of HMM on PHHM controlling for age and education.

## Discussion

Overall the most important explaining factor (in terms of total variance explained) for being in an ancestral homogamous vs. heterogamous marriage (HHM), for number of children, as well as for childlessness is the ancestral background: this complies with 31.4% of the variance explained in HHM, 14.1% of the variance in childlessness and 12.1% of the variance in the number of children. Hence ancestry is the most important predictor for ancestral homogamous vs. heterogamous marriages in the US. A limitation is that we do not know how far the ancestry dates back and we can only assume that the longer migration dates back, the weaker the effects of ancestry should be. To ascertain that not only the first generation migrants may have been responsible for the found associations we conducted the same analysis excluding all individuals not born in the US. The principal patterns of estimates, significances and variance explained remained unchanged, indicating that “ancestry effects” are universal and not caused be first generation migrants only (see supplement Table S2-S3).

Apart from ancestry, the ratio of an ancestry group in a county explains most of the variance in HHM (6%) - the positive association indicates, that the more individuals of a certain ancestry live in a county the more individuals marry within their own ancestry group. Thus, if more individuals of their own ancestry are at hand for marriage, individuals tend to marry homogamously, a pattern that was previously reported by Blau (1982) and Thomas (1951). recently, we have also found comparable marriage patterns on the basis of religious denominations in Europe (Fieder et al. 2019).

Apart from ancestry, second most of the variance (~1.25%) in HHM is explained by education: the higher the level of education of a woman, the higher is the probability of a homogamous marriage. Her spouse’s income in contrast, is significantly negatively associated with being in a homogamous marriage. Presumably, this is a result of female mate choice in favour of a wealthy spouse so that spousal income may become more important and thus relaxes mate choice based on ancestry. Interestingly, also the age of first marriage is negatively associated with ancestral homogamy, indicating that the older a woman at her marriage, the higher the probability that she marries outside her ancestry.

Ancestry also explains with 12% most of the variance of a woman’s number of children. Although HHM is significantly positively associated with the number of children, it explains only 0.07% of the variance. Out of the fixed factors, age of first marriage explains most of the variance (8.7%) and it is significantly negatively associated with number of children: the later a women marries the fewer children she has; a phenomenon well known as “postponing” (Schmidt 2012). The age of first marriage is thus a strong predictor for a woman’s number of children.

The patterns of associations of number iof children and SES-indicators were as expected: a woman’s income is significantly negatively but her husband’s income is significantly positively associated with number of children. Education has no significant association with number of children, which may be explained by the fact that education leads to a postponing of age at first marriage, which then appears to become the dominating variable in the model. This effect becomes further evident in the model of childlessness: higher education is associated with a lower probability of remaining childless as age of first marriage is included in the model. But again from the fixed factors age of first marriage explains most of the variance (12%). HHM is significantly positive associated with having at least one child, but explains only 0.09% of the variance in having at least one child. Women’ own income is significantly negative and spouse’ income positive associated with having at least one child. However as in the case of number of children age of first marriage conjoins a lot of the variance. The ratio of an ancestry group in a county is in no significant association to childlessness.

Albeit the effects of HHM on both, the number of children and childlessness are small in terms of total variance explained, these effects exist whether or not first generation migrants are included in the model. Thus ancestry even from an ancestral rather similar background (European ancestry), seems to influence the fertility of US women. This is to some extend comparable to Fox (2015) and Helgason (2008), demonstrating, that marrying a genetically more distant individual is associated with a loss in fertility. But on the other hand, marrying a genetically too closely related individual is associated with the risk of inbreeding (reviewed in Schahbasi et al. in press). Additionally, we aimed to estimate the “heritability” of marrying homogamously vs. heterogamously using a generalized linear mixed model https://cran.r-project.org/web/packages/QGglmm/vignettes/QGglmmHowTo.pdf) (de Villemereuil et al. 2016). We found a rather low heritability of 11.18%. However, a limitation of our approach is that we are not able to distinguish between additive genetic effects and the effects of the parental home (common environment), thus our estimate includes both additive genetic effects and the “common environment” (nature and nurture). Nonethelesse, to our knowledge, we are the first who ever tried to use a glmm on demographic family data to estimate “heritability”. This may be a promising approach in future studies to complement analyses on the “phenotypic level”.

On basis of our data we are not able to distinguish between genetic and social inheritance of the predisposition of ancestral homo- or heterogamy. However, in accordance with the first law of behavioural genetics - “that all human behavioural traits are heritable” - we assume that also the tendency to marry within or outside the ancestral group should have a genetic predisposition (Turkheimer 2000). Although with a total value of 11.8% variance explained by genetic and social heritability, only a small heritability is indicated. Nevertheless, according to Fischer (1930) and Falconer (1960), also traits with a comparable low heritability can be strongly selected. Concerning selection it is important to keep in mind, that homogamy per se does not change allele frequencies, but if there is an increase in the frequency of homozygotous individuals, it does provide a basis on which selection may act, for instance via selection against the recessive homozygote (Relethford 2012). Moreover, if the tendency to marry someone similar on one or more traits, has a genetic basis and assortative mating on this/these traits may lead to an increase in reproduction, a predisposition of assortative mating will spread in a population.

We conclude, that albeit only individuals of European ancestry have been included in the analysis (and irrespective whether or not first generation migrants are included) ancestry still has a relevant impact on marital and reproductive behavior. For women, the aviability of a potential spouse of the according ancestral group influences if they marry within or outside of their group. Furthermore, the tendency of marrying within or outside the own ancestral group may also have a heritable (genetic and/or social) component and thus selection may have been acting on this trait. From a methodological perspective we hope to encourage the analysis of heritability on the basis of large human data-sets such as census data sets in the future.

## Acknoweldegments

*IPUMS USA, University of Minnesota, www.ipums.org*.

## Data accessibility

All data sets used in the manuscript were provided by IPUMS USA: https://usa.ipums.org/usa/.

## Competing Interest

The authors declare no competing interests.

## Author contributions

A.S. interpreted the data and wrote the manuscript. S. H. interpreted the data and wrote the manuscript. M. F. conducted the analyses, interpreted the data and wrote the manuscript. All authors read and approved the final manuscript.

## Research Ethics

This article does not present research with ethical considerations.

## Notes

### Competing Interest Statement

The authors have declared no competing interest.

