## Supplementary Tables for "Marriage in the Melting Pot: An evolutionary approach to European ancestry, homogamy and fertility in the United States"

**Supplement**

| **code** | **education** |
|---|---|
| 0.00 | N/A (or None, 1980) |
| 1.00 | None |
| 2.00 | Nursery school |
| 3.00 | Kindergarten |
| | Elementary school: |
| 4.00 | 1st grade |
| 5.00 | 2nd grade |
| 6.00 | 3rd grade |
| 7.00 | 4th grade |
| 8.00 | 5th grade |
| 9.00 | 6th grade |
| 10.00 | 7th grade |
| 11.00 | 8th grade |
| | High school: |
| 12.00 | 9th grade |
| 13.00 | 10th grade |
| 14.00 | 11th grade |
| 15.00 | 12th grade |
| | College: |
| 16.00 | 1st year |
| 17.00 | 2nd year |
| 18.00 | 3rd year |
| 19.00 | 4th year |

| | |
|---|---|
| 20.00 | 5th year or more (40-50) |
| 21.00 | 6th year or more (60,70) |

Table S1) Encoding of highest education.

| | Beta Values | Std.Error | t-value | p-value | % Variance |
|---|---|---|---|---|---|
| (Intercept) | 0.000 | 0.145 | -7.915 | 0.000 | |
| AGE | 0.041 | 0.001 | 5.373 | 0.000 | 0.032 |
| AGEMARR | -0.063 | 0.001 | -8.163 | 0.000 | 0.111 |
| EDUC | -0.368 | 0.002 | -43.267 | 0.000 | 1.063 |
| INCTotal_OnlyPositive | -0.011 | 0.000 | -1.448 | 0.148 | 0.067 |
| INCTotal_OnlyPositive_SP | -0.093 | 0.000 | -11.503 | 0.000 | 0.319 |
| Ratio_Ancestry_Group_County | 1.047 | 0.075 | 62.660 | 0.000 | 6.450 |
| DF | 342786 | | | | |
| (Intercept) | Residual | | | | |
| StdDev: | | | | | |
| Variance expl. random | 13.6 | | | | |

Table S2) General linear mixed model of, age, age at first marriage, education, income, income of the spouse, the ratio of the own ancestry group in a county, regressing on HHM on basis of a binomial error structure with ancestry as random factor; excluding all individuals not born in the US.

| | Beta Values | Std.Error | t-value | p-value | % Variance |
|---|---|---|---|---|---|
| (Intercept) | 0.000 | 0.020 | 113.647 | 0.000 | |
| age | -0.019 | 0.000 | -33.662 | 0.000 | 0.821 |

| | | | | | |
|---|---|---|---|---|---|
| age marriage | -0.090 | 0.000 | -129.326 | 0.000 | 8.190 |
| ancestry same (ref. no) | 0.007 | 0.002 | 12.480 | 0.000 | 0.095 |
| education | 0.001 | 0.001 | 1.185 | 0.236 | 0.429 |
| total income | -0.028 | 0.000 | -45.059 | 0.000 | 1.064 |
| total income spouse | 0.008 | 0.000 | 13.183 | 0.000 | 0.025 |
| ratio ancestry county | -0.004 | 0.022 | -3.311 | 0.001 | 0.001 |
| DF | 342785 | | | | |
| (Intercept) | Residual | | | | |
| StdDev: 0.09847675 | 1.050 | | | | |
| Variance expl. random | 0.1 | | | | |

Table S3) General linear mixed model of, age, age at first marriage, education, income, income of the spouse, the ratio of the own ancestry group in a county and HHM, regressing on number of children on basis of a poison error structure with ancestry as random factor; excluding all individuals not born in the US.

| | **Beta Values** | **Std.Error** | **t-value** | **p-value** | **% Variance** |
|---|---|---|---|---|---|
| (Intercept) | 0.000 | 0.094 | 66.576 | 0.000 | |
| age | -0.238 | 0.002 | -9.423 | 0.000 | 0.904 |
| age marriage | -2.477 | 0.001 | -132.502 | 0.000 | 12.104 |
| ancestry same (ref. | 0.170 | 0.014 | 6.432 | 0.000 | 0.098 |

| | | | | | |
|---|---|---|---|---|---|
| no) | | | | | |
| education | 0.233 | 0.003 | 8.683 | 0.000 | 0.102 |
| total income | -0.624 | 0.000 | -28.791 | 0.000 | 1.007 |
| total income spouse | 0.590 | 0.000 | 20.557 | 0.000 | 0.888 |
| ratio ancestry county | 0.025 | 0.123 | 0.495 | 0.620 | 0.032 |
| DF | 342785 | | | | |
| StdDev: 0.09062322 | 0.997 | | | | |
| Variance expl. random | 13.600 | | | | |

Table S4) General linear mixed model of HHM age, age at first marriage, education, income, income of the spouse, the ratio of the own ancestry group in a county, regressing childlessness on basis of a binomial error structure with ancestry as random factor; excluding all individuals not born in the US.
